# Pharmacological inhibition of mitochondrial fission mitigates experimental thoracic aortic aneurysm

**DOI:** 10.1101/2025.03.23.644843

**Authors:** Anshul S. Jadli, Karina P. Gomes, Megan B. Meechem, Cameron D.A. Mackay, Noura N. Ballasy, Darrell Belke, Paul W.M. Fedak, Vaibhav B. Patel

## Abstract

Thoracic aortic aneurysm (TAA) is increasingly recognized as a vascular degenerative disease and is typically diagnosed incidentally due to its asymptomatic nature. TAA is clinically associated with the dilation of the aorta and excessive vascular remodeling, which can lead to aortic rupture if left untreated. Angiotensin II (Ang II), an active peptide in the renin-angiotensin system, has been widely implicated in the development of TAA. The current study investigated the role of mitochondrial dynamics in the onset and progression of TAA.

Male ApoEKO mice (8-10 weeks old) were infused with Ang II (1.44 mg/kg/day) and treated with mitochondrial division inhibitor 1 (mdivi-1) at a dose of 0.12 mg/kg/day. After 4 weeks of Ang II infusion, the ApoEKO mice developed TAA. Excessive mitochondrial fission was observed in thoracic vascular smooth muscle cells (vSMCs) using transmission electron microscopy (TEM) and confocal microscopy in response to angiotensin II (Ang II) treatment, which was reduced by mdivi-1. Echocardiographic and histological analyses of the Ang II-infused thoracic aorta revealed increased aortic dilation, vascular remodeling, and perivascular fibrosis. Pathological changes associated with Ang II-induced TAA were mitigated by mdivi-1. Co-treatment of isolated adult murine thoracic aortic vSMCs with mdivi-1 resulted in decreased mitochondrial fission, hyperproliferation, and phenotypic switching, accompanied by improved mitochondrial metabolism. Treatment with mdivi-1 inhibited Ang II-induced NF-κB signaling in vSMCs by blocking the nuclear translocation of p65 and cell cycle markers, thereby limiting inflammation and hyperproliferation.

In conclusion, the inhibition of mitochondrial fission reduced pathological changes by mitigating vascular remodeling, vSMC hyperproliferation, and vSMC phenotypic switching associated with Ang II-induced TAA. The clinical utility of inhibiting mitochondrial fission warrants further investigation, which may lead to a novel therapeutic strategy for preventing TAA development.

## Introduction

Thoracic aortic aneurysm (TAA) is a localized, degenerative, and permanent vascular disease characterized by a dilatation of the aorta exceeding 50% of its normal diameter. Often diagnosed incidentally because of its asymptomatic nature, TAA carries a mortality rate of approximately 70% among untreated patients (1, 2). Clinically, TAA is linked to aortic dilation and significant vascular remodeling, which includes the degradation of elastin fibers and the deposition of collagen around blood vessels (1, 2). Because various environmental and genetic factors contribute to the pathogenesis of TAA, the exact molecular mechanisms involved in TAA development remain unclear.

The renin-angiotensin-aldosterone system (RAAS) plays a crucial role in regulating blood pressure and maintaining vascular homeostasis, but its overactivity contributes to the development of TAA. Angiotensin II (Ang II), a primary effector of RAAS, promotes vascular inflammation, oxidative stress, endothelial dysfunction, and fibrosis mainly through Ang II type 1 (AT1R) receptors and, to a lesser extent, Ang II type 2 (AT2R) receptors (3, 4). Ang II plays a role in TAA pathogenesis by triggering hypertension, immune cell infiltration, extracellular matrix (ECM) degradation, and vascular remodeling (5, 6). In murine models, Ang II infusion causes TAA with pathological features akin to human disease, including aortic wall thickening, elastin fragmentation, luminal dilation, and macrophage accumulation (7, 8) Considering the translational relevance of this model, it has been extensively utilized to explore TAA mechanisms and possible therapeutic strategies.

Treatments for chronic aortic aneurysms include drug therapy and surgery for aneurysms that have a substantial risk of rupture. Medications such as beta-blockers, angiotensin receptor antagonists, doxycycline (a matrix metalloproteinase [MMP] inhibitor), anti-inflammatory agents (cyclooxygenase inhibitors), statins, and immunosuppressants (rapamycin) have been tested in animal models or clinical trials (9–12). However, none of these pharmacological methods has been effective in reducing the progression or risk of rupture of aortic aneurysms. Surgical options include surgical repair, which entails inserting an intraluminal graft through open access to the aneurysmal aorta or endovascular stenting (13). These surgical procedures carry a significant perioperative mortality risk and are, therefore, reserved for high-risk aneurysms. The lack of clinically relevant therapeutic interventions for aortic aneurysms stems from the complex pathophysiology related to the initiation and progression of TAA.

vSMCs are the primary cell type in the arterial media and play a crucial role in maintaining vascular tone. vSMCs exhibit phenotypic plasticity and, in response to pathological stimuli, can transition from a contractile to a synthetic phenotype (14). This phenomenon is known as vSMCs phenotypic switching and has been implicated in the development of various vascular diseases, including atherosclerosis and pulmonary hypertension (15–17). Recently, mitochondrial dynamics—encompassing fission and fusion— have been recognized as key factors in the phenotypic switching, proliferation, and migration of vSMCs (18–20). Dynamin-related protein 1 (Drp1) is a crucial regulator of mitochondrial fission and plays a role in maintaining mitochondrial homeostasis. Impaired regulation of mitochondrial fission has been linked to cardiovascular diseases such as ischemic/hypoxic injury, hypertension, and atherosclerosis (14, 21, 22). Drp1 has been documented as a primary contributor to excessive pathological mitochondrial fission, known as mitochondrial fragmentation. Given that excessive mitochondrial fission has been increasingly reported in vascular diseases, inhibiting Drp1 activity has been proposed as a potential target for treatment development.

Mitochondrial division inhibitor 1 (mdivi-1) is a quinazolinone that inhibits mitochondrial fission by reducing Drp1 GTPase activity (23). Mdivi-1 has demonstrated therapeutic effects in various conditions, including neurodegenerative diseases, acute myocardial infarction, hypertension, and sepsis (14, 24–26). A recent study showed the positive effects of inhibiting mitochondrial fission in hypertension by reducing Ang II-mediated phenotypic switching in vSMCs (14). These studies suggested a link between inhibiting mitochondrial fission, the phenotype of vSMCs, and the development of vascular disease. In this study, we examined the impact of mdivi-1 treatment on the pathogenic mechanism of Ang II-induced TAA in an experimental murine model.

## Materials and methods

### Experimental animals

*ApoE*-deficient mice (ApoEKO; C57BL/6 background) were obtained from the Jackson Laboratory (Bar Harbor, ME). The mice were kept in pathogen-free conditions and had access to sterilized food and water ad libitum. All animal experiments were conducted at the Department of Physiology and Pharmacology at the Cumming School of Medicine, University of Calgary. Alzet micro-osmotic pumps (Model 1004, Durect Corp.) were implanted subcutaneously in 8- to 10-week-old male mice to deliver Ang II (1.5 mg/kg/day) or saline (control) for 4 weeks (27). A subgroup of ApoEKO mice with Ang II osmotic minipumps was injected intraperitoneally with mdivi-1 (1.2 mg/kg) daily after surgery. All experiments were conducted in accordance with the guidelines of the University of Calgary Animal Care and Use Committee and the Canadian Council of Animal Care.

### Transthoracic echocardiography

Ultrasonic images of the aorta were acquired using a Vevo 3100 high-resolution imaging system, which features a real-time microvisualization scan head (MX550D, Visual Sonics, Toronto, Canada), in mice that were anesthetized with 2% isoflurane (500 ml/min oxygen flow) (27, 28). The aortic diameters at the thoracic aorta were measured using M-mode. The maximum aortic lumen diameter (systolic diameter corresponding to cardiac systole) and the minimum aortic lumen diameter (diastolic diameter corresponding to cardiac diastole) were monitored through simultaneous ECG recordings and used to calculate the aortic expansion index [(Systolic aortic diameter - Diastolic aortic diameter)/Systolic diameter X 100]. The VEVO vascular package’s (Vevo LAB software version 5.8.0) B-mode and EKV images determined the aortic distensibility.

### Isolation and culture of vSMCs

Vascular smooth muscle cells (vSMCs) were isolated from thoracic aortas obtained from 8- to 10-week-old ApoEKO mice and cultured as previously described (27, 29). Briefly, aortas were harvested under aseptic conditions, and the periaortic adipose tissues were promptly removed. The thoracic aortas were incubated with a digestive enzyme solution containing 1 mg/mL collagenase II (#LS004174, Worthington Biochemical) and 1 mg/mL elastase (#LS002292, Worthington Biochemical), supplemented with 1% penicillin/streptomycin (#CA12001-712, VWR) at 37°C in a 5% CO2 incubator for 5 minutes. The adventitial and intimal layers of the aorta were then removed using a surgical microscope. The medial layers of the aortas were cut into small pieces and incubated with the digestive enzyme solution for 90 minutes at 37°C in a 5% CO2 incubator. After digestion, the cells were washed and cultured in a sterile DMEM/F12 cell culture medium (#11330057, Gibco) containing 20% fetal bovine serum (FBS, #26140079, Gibco), supplemented with 1% penicillin/streptomycin (#CA12001-712, VWR). The cells were passaged upon reaching 90-100% confluency. Thoracic aortic vSMCs were serum-deprived for 24 hours by incubation in DMEM/F12 medium with 1% fetal bovine serum and penicillin/streptomycin. The cells were challenged with Ang II (100 nM; #05-23-0101, Millipore Sigma) in DMEM/F12 for 24 hours. In the treatment group, mdivi-1 (50 μM; #S7162, Selleck Chemicals, USA) was added to the vSMCs 30 minutes before co-treatment with Ang II for 24 hours. A subgroup of vSMCs without Ang II and mdivi-1 treatment, treated with saline, served as controls.

### Transmission electron microscopy

For transmission electron microscope analysis, samples of mouse aorta tissues were fixed in 2.5% glutaraldehyde, then washed three times with phosphate-buffered saline, followed by post-fixation in 1% osmium tetroxide for 1 hour. The samples were stained with 2% aqueous uranyl acetate and dehydrated in a graded series of ethanol. After infiltration and polymerization, ultrathin sections were cut, flattened with xylene vapor, collected on nickel grids, and observed using a Hitachi H-7650 transmission electron microscope (Hitachi, Japan) at magnifications of 10000x and 25000x. Quantitative measurements of mitochondrial morphology were performed using the mitomorphology plugin in Fiji/ImageJ (version 1.52i) software.

### Histology and immunofluorescence staining

Mice were perfusion-fixed at 80 mmHg with buffered formalin after 4 weeks of Ang II (±mdivi-1) or saline infusion, which preserved the blood vessels in their native state, as previously described (27). Subsequently, aortas were collected and fixed in 10% buffered formalin for 48 h. Five μm thick formalin-fixed paraffin-embedded (FFPE) sections were utilized for histological staining, including Hematoxylin and Eosin (H&E), Gomori trichrome, and Verhoeff Van Gieson (VVG), as previously described (27). Picrosirius red (PSR) staining was performed following established protocols (30). Collagen-positive regions in the thoracic aorta were imaged using a confocal microscope (Leica SP8) and quantified through morphometric analysis with Fiji/ImageJ software (version 1.52i). Additionally, five μm thick FFPE sections were used for immunofluorescence staining, as previously described (27). After deparaffinization, sections were immersed in EDTA-citrate buffer (pH 6.2) for antigen retrieval. Subsequently, sections were washed with PBS and permeabilized with 0.1% Triton X-100, followed by blocking with 4.0% bovine serum albumin (BSA). Sections were incubated overnight at 4°C with the corresponding primary antibodies for α-smooth muscle actin (α-SMA, 1:500, #ab32575, Abcam), followed by incubation with secondary antibodies for Alexa Fluor 488-conjugated goat anti-rabbit (1:1000, #A11034, Invitrogen) at room temperature for 1 hour. Prolong Gold with DAPI (#P36935, Invitrogen) served as a mounting medium and nuclear stain. Immunostained aorta sections were imaged using a line-scanning confocal microscope (Leica SP8) and analyzed with Fiji/ImageJ software (version 1.52i).

After 24 hours, saline, AngII, and mdivi-1 treated vSMCs were fixed with 4% paraformaldehyde for 15 minutes, then washed with PBS three times. Fixed thoracic vSMCs were then permeabilized for 10 minutes with 0.1% Triton X-100 at 4°C. The cells were treated with 4% BSA for 1 hour at room temperature and then incubated with a primary antibody for NF-kB p65 (1:250, #8242, Cell Signaling) overnight at 4°C, followed by treatment with secondary antibodies for Alexa Fluor 488-conjugated goat anti-rabbit (1:1000, #A11034, Invitrogen) at room temperature for 1 hour. ProLong Gold with DAPI (#P36935, Invitrogen) was used as the mounting medium and for nuclear staining. Cells were imaged using a confocal microscope (SP8, Leica) and analyzed with Fiji/ImageJ (version 1.52i) software.

### Assessment of mitochondrial dynamics

The mitochondrial structure was evaluated using MitoTracker Red CMXRos (#M7512, Invitrogen) staining and confocal imaging. Briefly, control saline or Ang II (±mdivi-1)-treated cells were incubated with 100 nM MitoTracker Red CMXRos for 30 minutes at 37°C in a CO2 incubator. MitoTracker Red-labeled cells were fixed with 4% paraformaldehyde and imaged with the Leica SP8 confocal microscope. Mitochondrial fragmentation was quantified using Fiji/ImageJ (version 1.52i) and the Mitochondrial Network Analysis (MiNA) plugin (31).

### Mitochondrial bioenergetics

The mitochondrial bioenergetic profile was determined using the Seahorse Analyzer XFe24 (Agilent Technologies). Thoracic vSMCs were plated at 40000 cells per well on XFe24 plates. After treatment with either vehicle or Ang II, with or without mdivi-1, the cells were switched to XF media (non-buffered DMEM containing 2 mM L-glutamine, 100 mM sodium pyruvate, and 5 mM glucose) 30 minutes before measuring bioenergetic parameters. The oxygen consumption rate (OCR) and extracellular acidification rate (ECAR) were measured using the XFe24 Cell Mito Stress Test Kit (#103015-100, Agilent Technologies). Baseline and stressed OCR and ECAR were determined prior to and following the injection of stressor compounds: 1 μM oligomycin (an ATP synthase inhibitor), 1 μM fluorocarbonyl cyanide phenylhydrazone (FCCP, an uncoupler), 1 μM rotenone (a Complex I inhibitor), and 1 μM antimycin A (a Complex III inhibitor). The data were analyzed using Wave software version 2.6 (Agilent Technologies) and presented as the metabolic potential for OCR and ECAR.

### Assessment of vSMC proliferation

Thoracic aortic SMC proliferation was assessed using Ki67 staining and flow cytometric analysis as previously described (32). Briefly, vSMCs were plated at 25000 cells per well in 6-well plates and treated with vehicle (saline) and Ang II (±mdivi-1) for 24 hours. The saline and Ang II (±mdivi-1)-treated cells were trypsinized, and the monolayer cells in the cell culture medium were centrifuged at 1500 x g for 5 minutes. The cell pellets were fixed with 70% ethanol and incubated at -20°C for 2 hours. After fixation, the cells were washed twice with staining buffer (1% FBS, 0.09% NaN3 in PBS) and resuspended in staining buffer at 1 × 10^6 cells/mL. The cells were then incubated with the Anti-Ki67 antibody (#ab16667, Abcam) at room temperature for 30 minutes. Following incubation, the cells were washed twice with staining buffer. Alexa Fluor 488-conjugated goat anti-rabbit secondary antibody (#A11034, Invitrogen) was added, and the cells were incubated at room temperature for 30 minutes in the dark. After incubation, the cells were washed twice with staining buffer. Samples were analyzed within 1 hour using the Attune NxT flow cytometer (ThermoFisher Scientific, MA, USA) at 488 nm excitation and 530 nm emission wavelengths. Quantitative analysis of the flow cytometric data was performed using Invitrogen™ Attune™ NxT Software (version 2.6, ThermoFisher Scientific, MA, USA).

### Quantitative real-time PCR

Total RNA was extracted from thoracic aortic vSMCs using the TRIzol reagent (#15-596-018, Thermo Fisher Scientific) following the manufacturer’s guidelines. The isolated RNA was dissolved in RNase-free water, and its concentration was measured with a spectrophotometer. cDNA was subsequently prepared using the cDNA Reverse Transcription Kit (#4374966, Thermo Fisher Scientific). TaqMan probes for quantitative PCR were then utilized to assess the expression of specific genes: Acta2 (Mm.PT.58.16320644), Cnn1 (Mm.PT.58.12652862), Myh11 (Mm.PT.58.9236105), Il-6 (Mm.PT.58.10005566), Mmp2 (Mm00439498_m1), Mmp9 (Mm00442991_m1), Col1a1 (Mm00801666_g1), Col3a1 (Mm00802331_m1), ccne1 (Mm.PT.58.29564054), ccnd1 (Mm.PT.58.28503828), Cdkn1a (Mm.PT.58.17125846), Chuk (Mm.PT.58.13005707), Dnm1l (Mm.PT.58.32328364), and 18S rRNA (Hs.PT.39a.22214856.g). For each gene, a standard curve was constructed using known concentrations of cDNA (0.1, 1, 10, 100, and 1000 ng) as a function of cycle threshold (CT). Expression analysis of the indicated genes was conducted using TaqMan Real-time PCR on the QuantStudio™5 system (Thermo Fisher Scientific, MA, USA). Data were processed using the QuantStudio™ design and analysis software version 1.4.3. All samples were analyzed in triplicate in 384-well plates. mRNA expression data were presented as relative expressions to 18S rRNA, serving as an endogenous control.

### Statistical analysis

All data are presented as mean ± SEM. All statistical analyses were conducted using GraphPad Prism v9 (San Diego, CA). A one-way ANOVA followed by Tukey’s multiple comparisons post hoc analysis was employed to compare the data among the Saline, Ang II, and Ang II+mdivi-1 groups. A p-value of <0.05 was considered statistically significant.

## Results

### Ang II-mediated TAA potentiates mitochondrial structure damage and promotes fission

Excessive mitochondrial fission and structural damage mediated by Ang II have been reported in vascular pathologies (20, 33, 34). To assess the causal association between Ang II-mediated aortic aneurysm and mitochondrial morphology, we performed transmission electron microscopy (TEM) on the aneurysmal tissue (**Figure 1A**). TEM images exhibited extensive mitochondrial structural damage with mitochondrial fragmentation in the aortic tissue from Ang II-infused mice. ApoEKO mice treated with mdivi-1 showed mitochondria with preserved internal structural integrity comparable to the saline group. The quantification of morphometric parameters such as Feret’s diameter, form factor, perimeter, and surface area (**Figure 1B-E**) further corroborated the protective effects of mdivi-1 on Ang II-mediated mitochondrial damage in aortic tissues. Consistent with the TEM data, immunofluorescence assessment of mitochondrial structure using MitoTracker Red showed excessive fission in thoracic vSMCs treated with Ang II for 24 hours. Elongated mitochondria in vSMCs treated with saline or mdivi-1 suggested a reduction in mitochondrial fragmentation and preservation of structural integrity (**Figure 1F**). Quantification of MitoTracker Red-stained confocal images using MiNA (**Figure 1G**), an NIH ImageJ plugin, revealed decreased mean branch length and increased number of individuals, indicative of smaller mitochondria in Ang II-treated thoracic aortic vSMCs (**Figure 1H-M**). Treatment of thoracic aortic vSMCs with mdivi-1 resulted in elongated mitochondria and preservation of the number of individuals and mean branches per network, suggesting alleviation of Ang II-induced mitochondrial fission (**Figure 1H-M**).

**Figure 1:**
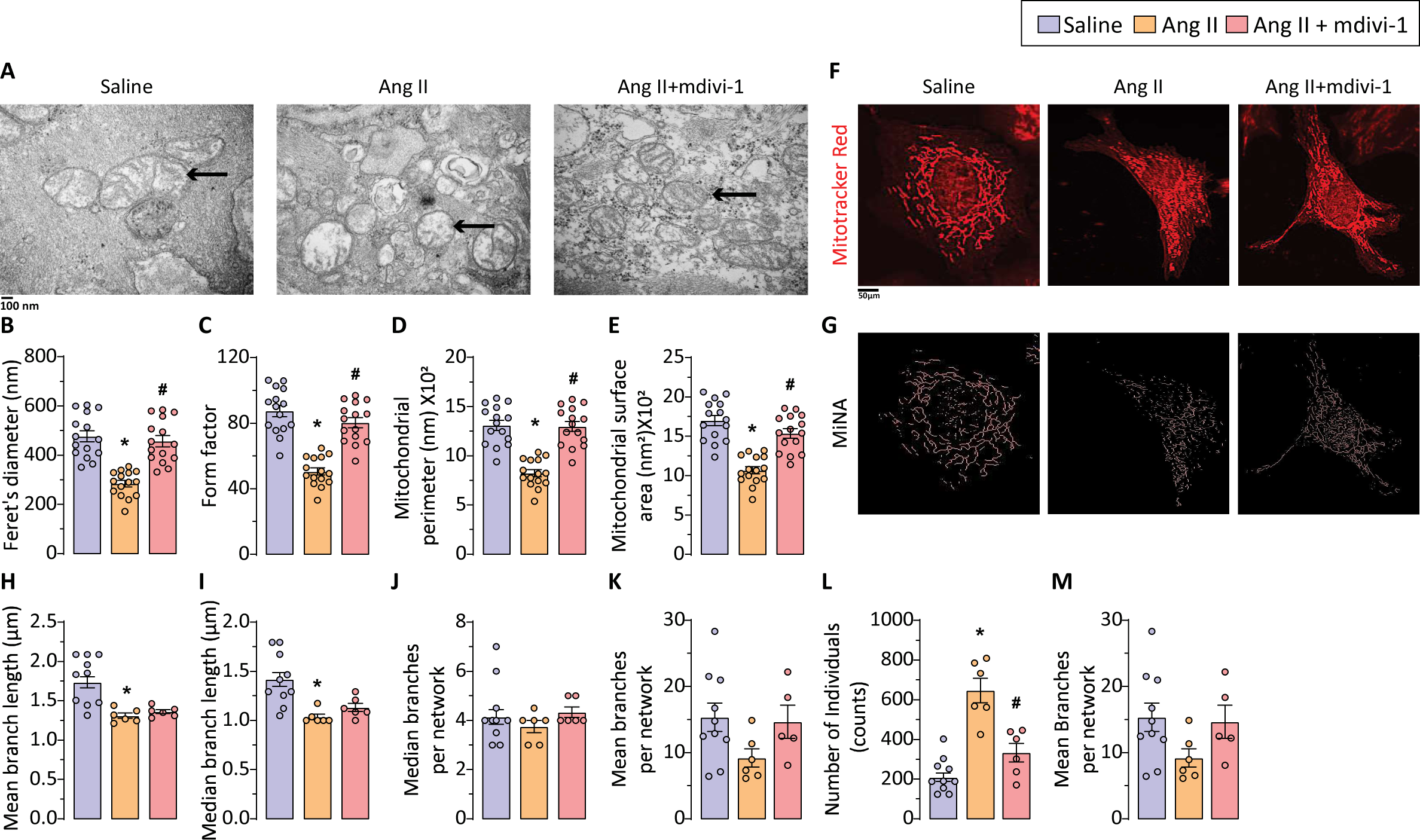
mdivi-1 treatment supresses Angiotensin II-mediated excessive fission and mitochondrial structural damage in thoracic aortic SMCs. Representative images from transmission electron microscope showing disruption in mitochondrial morphology in thoracic aortic SMCs from (A) *ApoEKO* mice aorta infused with Ang II for 4 weeks which was reversed by mdivi−1 treatment. The arrow indicates alteration in mitochondrial structure in aortic SMCs. Quantification of mitochondrial morphometric parameters in (B−E) mice using mito morphology plugin in ImageJ. Data points represent technical replicates from biological replicates (n = 3 in each group) (F) Confocal microscopy showed increased mitochondrial fragmentation using mitotracker red staining in thoracic SMC treated with Ang II (n=3). mdivi−1 (n=3) treatment prevented excessive mitochondrial fission which was quantified by (G) morphometric analysis of mitochondrial structure using mitochondrial network analysis (MiNA); (H−M) Quantification of mitochondrial structure parameters. Data points represent technical replicates from biological replicates (n = 5 in saline group) *, p < 0.05 compared with saline group; #, p < 0.05 compared with Ang II group using one−way ANOVA.

### Inhibition of mitochondrial fission prevents structural and functional alteration in the thoracic aorta

Infusing Angiotensin II into hypercholesterolemic ApoEKO mice is a well-established experimental model for TAA that stimulates various features of human TAA (8, 32). The Ang II-infused ApoEKO mice exhibited significantly higher mortality rates and a greater incidence of aortic rupture (**Figure 2A-B**). As shown previously (32), the 4-week infusion of Ang II into ApoEKO mice resulted in the development of TAA, as evidenced by increased aortic dilation. The aortic dilation, a key characteristic of aortic aneurysm, in Ang II-infused ApoEKO mice was measured using echocardiography. Representative echocardiography images and averaged aortic diameter showed significant aortic dilation in the thoracic aorta of Ang II-infused ApoEKO mice (**Figure 2C-H**). To investigate the protective effects of inhibiting mitochondrial fission, we administered mdivi-1 (1.2 mg/Kg/day) intraperitoneally to ApoEKO Ang II-infused mice. Treatment with mdivi-1 reduced pathological changes associated with Ang II-induced TAA (**Figure 2C-H**). The aortic expansion index, which indicates the elasticity of the aorta during systole and diastole, was not significantly decreased in the thoracic aorta of Ang II-infused mice (**Figure 2G**). Distensibility is commonly used to assess aortic stiffness and is defined as the aorta’s ability to expand during systole. The aortic distensibility in the thoracic aorta was measured using the high-frequency ECG-gated Kilohertz Visualization (EKV) mode of echocardiography. Representative ultrasound images displayed decreased distensibility of the thoracic aorta in Ang II-infused mice, which was restored with mdivi-1 treatment (**Figure 2H**). Altered ultrasound parameters confirmed the compromised structural and functional integrity of the aorta in Ang II-infused mice, which were reversed by the administration of mdivi-1. Thus, the mdivi-1-mediated reduction of Ang II-induced pathological structural and functional remodeling of the aorta suggests protective effects of pharmacological inhibition of mitochondrial fission in TAA.

**Figure 2:**
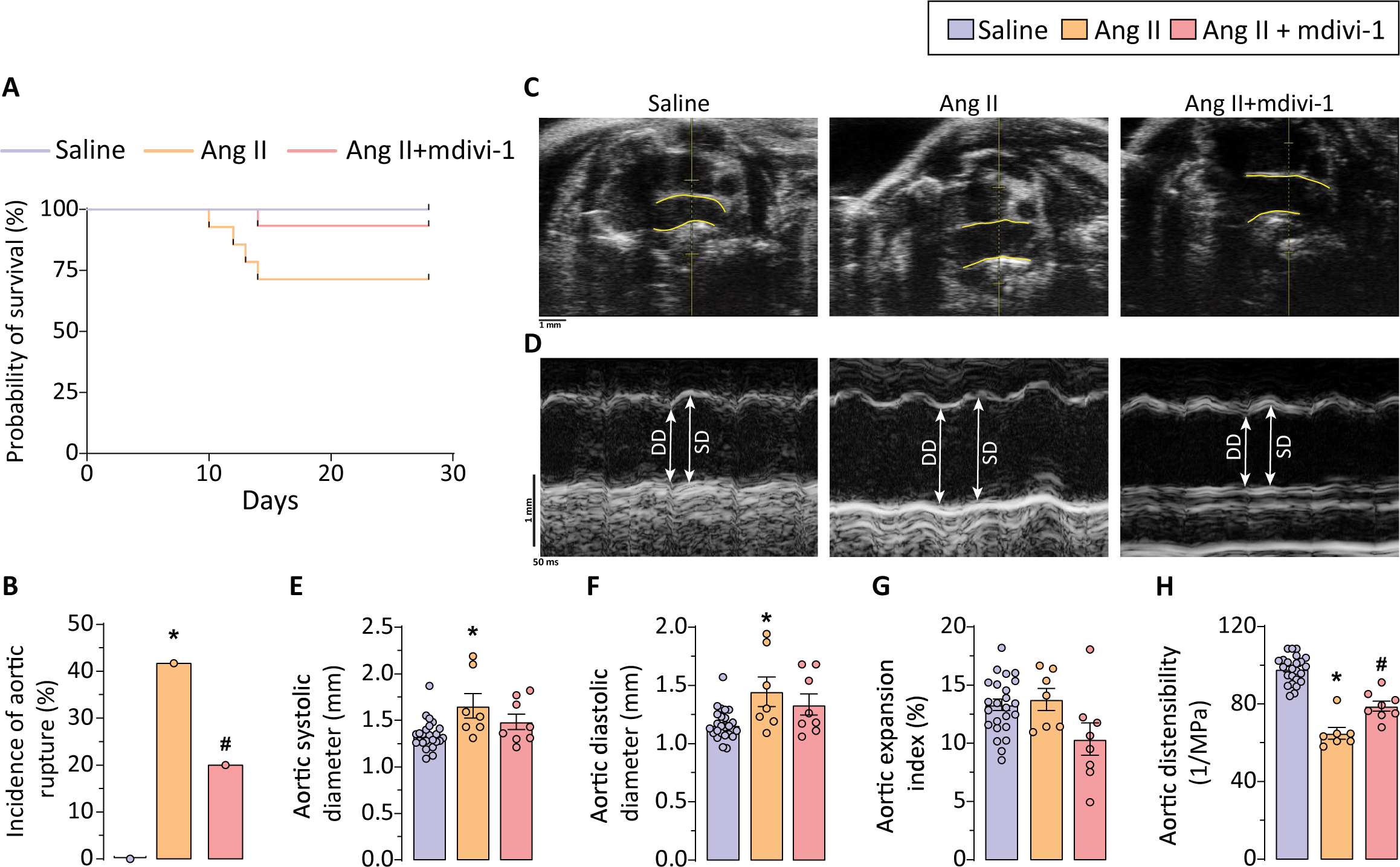
Inhibition of mitochondrial fission attenuates development of Ang II-induced thoracic aortic aneurysm in *ApoE* knockout mice. (A) The survival curve showed increased thoracic aortic rupture associated death in Ang II−infused mice (5/12 = 41.66%). Co−administration of mdivi−1 resulted in decreased mortality (2/10 = 20%) with (B) significantly reduced incidence of aortic rupture. (C) B−mode and (D) M−mode echocardiographic images of thoracic aorta and quantification of averaged aortic (E) systolic, (F) diastolic diameters, (G) aortic expansion index and (H) distensibility of thoracic aorta demonstrated increased aortic dilatation in response to Ang II (n=7) for 4 weeks and improvement in aortic diameter with midivi−1 treatment (n=8). Each data point on the graph represents a biological replicate. *, p < 0.05 compared with saline group (n=25); #, p < 0.05 compared with Ang II group using one-way ANOVA.

### Inhibition of excessive mitochondrial fission prevents Ang II-induced negative aortic remodeling

Previous studies have shown (32), the development of Ang II-induced TAA has been associated with extensive vascular remodeling. To examine the role of fission inhibition in mitigating the adverse aortic remodeling associated with TAA, we assessed key components related to the structural integrity of the aortic wall. Verhoeff-Van Gieson (VVG) staining revealed disrupted elastin fibers in the aorta of Ang II-infused mice; however, the administration of mdivi-1 restored the structural integrity of these fibers (**Figure 3A**). A clinical hallmark of TAA development in aneurysmal patients is decreased medial layer thickness, a condition we also observed in the thoracic aorta of Ang II-infused mice. Treatment with mdivi-1 resulted in reduced aortic wall degeneration and restored medial thickness in the thoracic aorta (**Figure 3B**).

**Figure 3:**
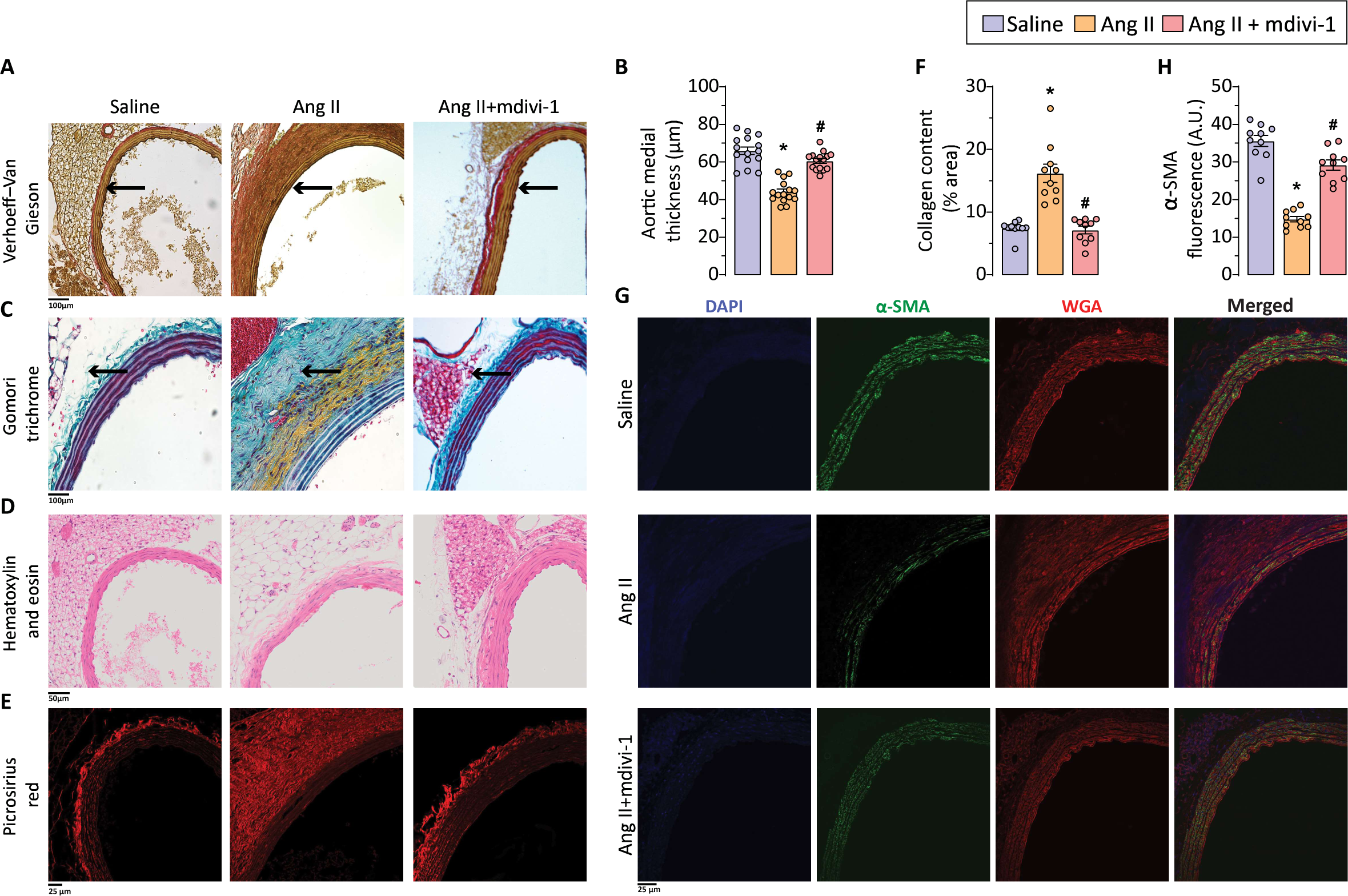
Suppression of excessive mitochondrial fragmentation attenuates Ang II-mediated adverse thoracic aortic remodelling. Representative images for (A) Verhoeff−Van Gieson (elastin); arrow indicates aortic medial layer, (C) Gomori trichrome (collagen); arrow indicates perivascular fibrosis, and (D) Hematoxylin and eosin (H&E) staining and quantification of (B) Aortic wall thickness exhibited reduced medial layer thickness in thoracic aorta with Ang II treatment. mdivi−1 administration restores structural integrity of aortic wall. Representative images for (E) Picrosirius red (collagen) staining and quantification of (F) collagen content (% area) showed excessive perivascular fibrosis with Ang II infusion which was reversed by mdivi−1 treatment. Representative images for (G) α−Smooth muscle actin (α−SMA) immunofluorescence staining and (H) quantification α−SMA fluorescence in thoracic aorta demonstrated reduced α−SMA expression in response to Ang II suggestive loss of contractile marker expression in thoracic SMCs. mdivi−1 reinforces contractile phenotype in thoracic SMCs. Data points represent technical replicates from biological replicates (n = 3 in each group) *, *p* < 0.05 compared with saline group; #, *p* < 0.05 compared with Ang II group using one−way ANOVA.

ECM remodeling, including excessive collagen deposition in the aortic adventitial layer, has been reported during the onset and progression of TAA (35, 36). Histological evaluation of collagen, a major component of the extracellular matrix, using Gomori trichrome staining revealed excessive collagen deposition in the adventitia of the Ang II-infused aorta, resulting in significant perivascular fibrosis (**Figure 3C**). Moreover, the loss of red coloration in the medial layer may indicate a loss of vSMC contractile phenotype and a transition to a synthetic phenotype. Excessive perivascular collagen deposition was also demonstrated by Picrosirius red staining, confocal imaging, and collagen content quantification in the thoracic aorta infused with Ang II (**Figure 3E-F**). Treatment with mdivi-1 significantly mitigated maladaptive aortic structural remodeling and perivascular fibrosis, leading to decreased collagen levels and deposition in the peri-adventitial region (**Figure 3E-F**). Histological analyses using Hematoxylin and eosin (H&E) staining showed the whitening of perivascular adipose tissue (PVAT) in the thoracic aorta of Ang II-infused mice (**Figure 3D**). Under physiological conditions, the PVAT surrounding the thoracic aorta displays brown adipose tissue-like characteristics (37), while pathological stimuli in Ang II-induced TAA cause adipose tissue dysfunction and whitening of adipocytes along with increased adipocyte hypertrophy (38, 39). The inhibition of mitochondrial fission via mdivi-1 treatment decreased PVAT remodeling and lessened adipocyte hypertrophy.

Alpha-smooth muscle actin (α-SMA) is a key contractile protein that forms the core of sarcomeres in vSMCs, and altered levels of α-SMA indicate a continuum of smooth muscle cell (SMC) phenotypes from contractile to synthetic. Immunofluorescence staining and confocal image analysis revealed a statistically significant decrease in α-SMA levels in the medial smooth muscle cells (**Figure 3G-H**), which further suggests a phenotypic switch of vSMCs to a synthetic phenotype in the Ang II infusion group. Mdivi-1 treatment restored α-SMA levels, highlighting the critical role of fission inhibition in reinforcing the contractile phenotype of thoracic vSMCs (**Figure 3G-H**). Our data also suggest that the protective effects of fission inhibition on Ang II-induced TAA may be, at least in part, mediated by mdivi-1’s effects on vSMCs.

### Inhibition of mitochondrial fission reduces Ang II-mediated detrimental effects on mitochondrial metabolism

As the Ang II infusion caused significant mitochondrial damage in the thoracic aorta, we assessed the impact of Ang II-induced mitochondrial damage on mitochondrial metabolism using the Seahorse Cell Mito Stress Test. Ang II treatment resulted in decreased mitochondrial respiration, as indicated by lower OCR and ECAR (**Figure 4A-B**). vSMCs demonstrated reduced ATP production, maximal respiration, spare respiratory capacity, and an increased proton leak following Ang II treatment (**Figure 4C-J**). Although the analysis did not reveal statistical significance, the inhibition of mitochondrial fission was associated with improved metabolic parameters. The negative marker of mitochondrial function, specifically the increase in proton leak, was heightened in thoracic vSMCs treated with Ang II and mdivi-1 treatment, which subsequently led to a reduction in proton leak. Maximal respiration capacity and spare respiratory capacity, which represent the cell’s bioenergetic reserve under stress conditions, were both enhanced following treatment with mdivi-1 (**Figure 4C-J**). Overall, this suggests that preventing mitochondrial fragmentation improved mitochondrial respiration.

**Figure 4:**
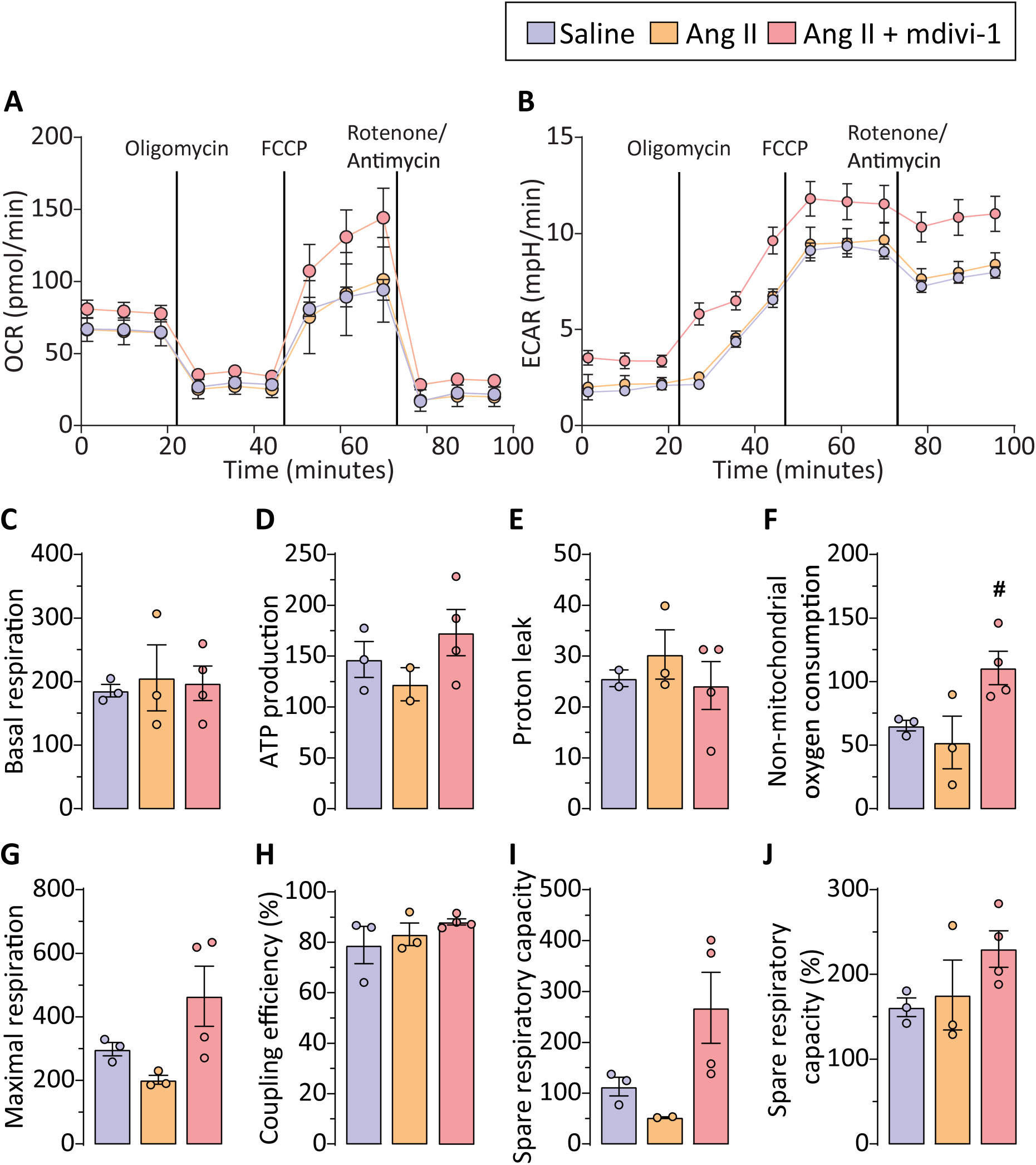
mdivi-1 treatment improves mitochondrial metabolism in thoracic aortic smooth muscle cells (SMCs). Analysis of mitochondrial metabolism using seahorse XFe24 showed altered mitochondrial metabolism in thoracic SMCs in response to Ang II treatment depicted by (A) oxygen consumption rate (OCR) and (B) extracellular acidification rate (ECAR) which was reversed by mdivi−1. (C−J) Quantification of metabolic parameters for mitochondrial metabolism did not show statistical significance but (D) ATP production, (E) proton leak, (F) Non−mitochondrial oxygen consumption, (G) maximal respiration and (I) spare respiratory capacity showed restoration of metabolic parameters with mdivi−1 (n=4) treatment indicating protective effect of mdivi−1 on thoracic SMCs via improved mitochondrial metabolism. Each data point on the graph represents a biological replicate. *, *p* < 0.05 compared with saline group (n=3); #, *p* < 0.05 compared with Ang II (n=3) group using one-way ANOVA.

### Inhibition of mitochondrial fission leads to reduced pathological phenotypic switching and hyperproliferation in vSMCs

The phenotypic plasticity of vSMCs is essential for vascular growth and remodeling. Most vSMCs display a contractile phenotype to regulate vascular tone. However, they can differentiate into a synthetic phenotype, which contributes to cardiovascular pathologies (40, 41). As previously shown, Ang II promotes phenotypic switching in vSMCs and contributes to the development of thoracic aortic aneurysm through matrix remodeling and inflammation (32). We assessed the impact of mdivi-1-mediated mitochondrial fission inhibition on the phenotypic switching of vSMCs derived from the thoracic aorta. The quantitative mRNA expression analysis revealed a decrease in the expression of contractile genes, including *Acta2*, *Cnn1*, and *Myh11* (**Figure 5A-C**) in Ang II-treated thoracic vSMCs. Furthermore, Ang II treatment also elevated the mRNA expressions of *Il-6*, *Mmp2*, *Mmp9*, *Col1a1*, and *Col3a1* (**Figure 5D-H**) in the vSMCs, indicating their polarization towards inflammatory and synthetic phenotypes. Notably, co-treatment with Mdivi-1 maintained the contractile phenotype of thoracic aortic vSMCs, as indicated by the preservation of mRNA expression of contractile genes (**Figure 5A-C**). Additionally, mdivi-1 treatment diminished the Ang II-induced onset of the inflammatory and synthetic phenotypes in thoracic aortic vSMCs, leading to reduced mRNA expression of *Il-6*, *Mmp2*, *Mmp9*, *Col1a1*, and *Col3a1* (**Figure 5D-H**). The data suggested that mdivi-1-mediated fission inhibition suppresses the polarization of thoracic vSMCs towards the synthetic phenotype.

**Figure 5:**
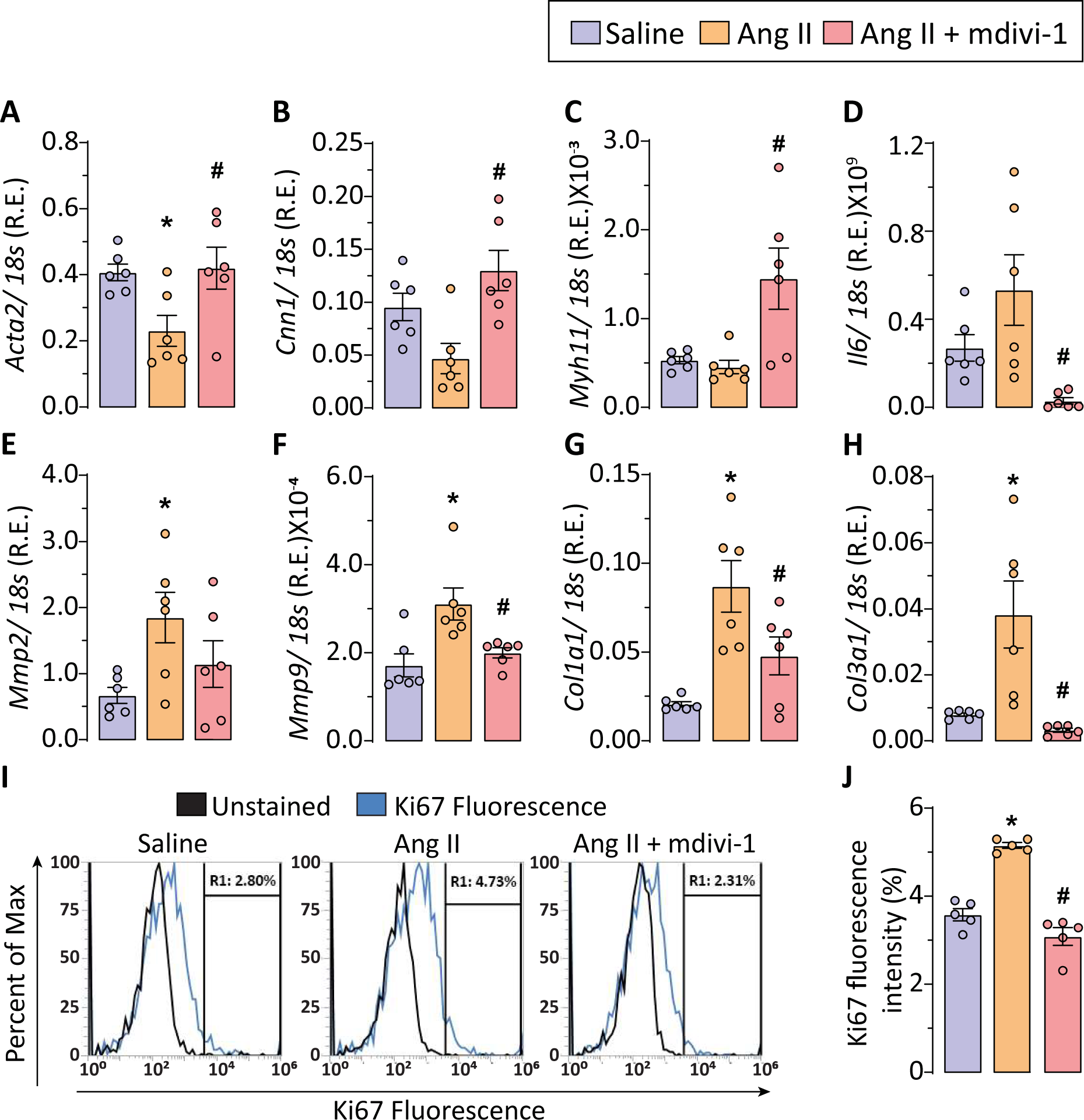
Mitochondrial fission inhibition supresses pathological phenotypic switching and hyper-proliferation in thoracic SMCs in response to Ang II. Gene expression profile for contractile SMCs markers i.e. (A) alpha−Smooth muscle actin (*Acta2*), (B) Calponin1 (*Cnn1*), (C) Myosin heavy chain 11 (*Myh11*) showed reduced expression in Ang II-treated thoracic SMCs. The inflammatory and synthetic markers i.e. (D) Interleukin−6 (*Il-6*), (E) Matrix metalloprotease 2 (*Mmp2*), (F) Matrix metalloprotease 9 (*Mmp9*), (G) Collagen I (*Col1a1*), and (H) Collagen III (*Col3a1*) showed increased expression in Ang II-treated SMCs (n=6). The gene expression for contractile and synthetic markers were normalized by mdivi−1 (n=6) treatment suggesting mitochondrial fission inhibition reinforces contractile phenotype in thoracic SMCs. (I) Flow cytometric analysis of Ki67 staining and (J) quantification of %Ki67 fluorescence indicated hyperproliferation in thoracic SMCs in response to Ang II (n=5) treatment which was normalized by mdivi−1 (n=5) treatment. Each data point on the graph represents a biological replicate. R.E.: Relative expression; *, p < 0.05 compared with saline group; #, p < 0.05 compared with Ang II group using one−way ANOVA.

As previously reported, Ang II encourages vSMC proliferation and migration, contributing to the development of heart failure, atherosclerosis, aortic aneurysm, and hypertension. (32, 42). *In vitro*, mRNA expression demonstrated that Ang II mediates phenotypic switching in thoracic vSMCs, and these dedifferentiated vSMCs reportedly exhibit increased cell proliferation (43). We used Ki67, a nuclear protein marker of active cell proliferation, to evaluate Ang II-induced thoracic vSMC proliferation. The flow cytometry analysis showed a significant increase in Ki67 fluorescence in thoracic vSMCs treated with Ang II (**Figure 5I-J**). Inhibition of mitochondrial fragmentation by mdivi-1 in vSMCs reduced Ki67 fluorescence, indicating decreased hyperproliferation in thoracic aortic vSMCs (**Figure 5I-J**). The data demonstrated Ang II-mediated phenotypic switching and hyperproliferation in thoracic vSMCs, contributing to vascular remodeling, which is attenuated by mdivi-1 treatment.

### Suppression of mitochondrial fission attenuates NF kB p65 signaling in thoracic vSMCs

The in vitro and in vivo experiments indicated that Ang II-induced hyperproliferation and phenotypic switching in thoracic vascular smooth muscle cells (vSMCs) play a causal role in the development of thoracic aortic aneurysm (TAA). Previously, Yoshida et al. reported that the inhibition of nuclear factor-kappa B (NF-κB) signaling, specifically in vSMCs, attenuates their phenotypic switching and proliferation (44). Furthermore, the role of mitochondrial dynamics in modulating NF-kB signaling in microglial cells highlights the interplay between Drp1-dependent fission and pro-inflammatory signaling (45). To understand the relationship between fission inhibition and the attenuation of phenotypic switching and hyperproliferation in thoracic vSMCs, we investigated the nuclear translocation of NF-κB p65, a subunit of NF-κB involved in transcriptional regulation. Thoracic vSMCs treated with Ang II exhibited nuclear translocation of NF-κB p65, accompanied by increased mitochondrial fission (**Figure 6A**). The inhibition of mitochondrial fission by mdivi-1 resulted in the reduction of nuclear translocation and cytosolic staining of NF-kB p65. Furthermore, the quantitative mRNA expression analysis demonstrated an increased expression of Ccne1 (Cyclin E1) and Ccnd1 in Ang II-treated thoracic VSMCs (**Figure 6B-C**). Cyclin E1 controls the G1-to-S phase transition, while Cyclin D1 regulates early G1 phase progression; both play critical roles in vascular smooth muscle cell proliferation and remodeling. The dysregulation of these genes has been linked to pathological vascular remodeling and hypertension-associated vascular diseases. (46, 47), Though the expression level was not statistically significant, Ang II treatment reduced the expression of *Cdkn1a*, cyclin-dependent kinase inhibitor 1/p21, in vSMCs (**Figure 6D**). The mdivi-1 treatment restored the expression of *Ccne1*, *Ccnd1,* and *Cdkn1a,* suggesting that inhibition of mitochondrial fission restores cell cycle regulation (**Figure 6B-D**). The data indicated that mdivi-1-mediated inhibition of Drp1-dependent fission attenuates NF-kB signaling, preventing p65 translocation, which consequently restores transcription of cell cycle proteins to prevent phenotypic switching and hyperproliferation in thoracic vSMCs.

**Figure 6:**
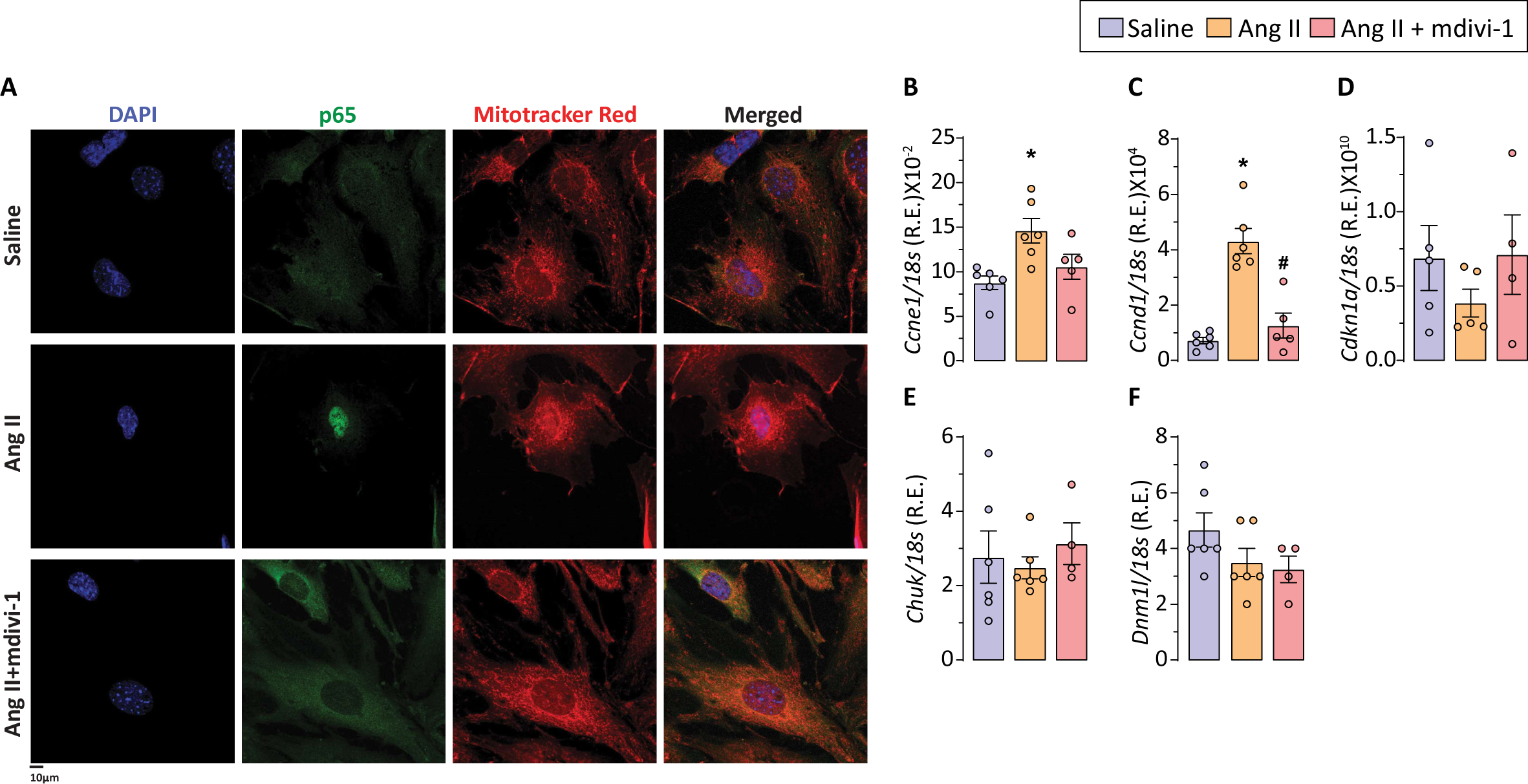
mdivi-1 attenuates nuclear translocation of NF kB p65 and cell cycle markers in thoracic SMCs. Representative images from confocal microscopy showed (A) increased nuclear translocation of p65 and mitochondrial fission in thoracic SMCs in response to Ang II treatment. mdivi−1 suppresses NF kB signalling by attenuating nuclear translocation of p65 subunit. Gene expression profile for cell cycle markers showed increased expression for (B) cyclin E1, (C) cyclin D1 and reduced expression for (D) cyclin−dependent kinase inhibitor p21 in thoracic SMCs treated with Ang II indicating higher cell proliferation rate. mdivi−1 treatment attenuated the SMCs hyperproliferation. The expression levels of (E) inhibitor of nuclear factor kappa−B kinase subunit alpha, Chuck and (F) Drp1 did not show significant difference between saline (n=6), Ang II (n=6) and Ang II+ mdivi−1 (n=5) groups. Each data point on the graph represents a biological replicate. R.E.: Relative expression; *, p < 0.05 compared with saline group; #, p < 0.05 compared with Ang II group using one−way ANOVA.

## Discussion

Thoracic aortic aneurysm (TAA) is a degenerative, irreversible dilatation of the thoracic aorta to at least 50% of the normal diameter. The incidence of TAA ranges from 5 to 10 cases per 100,000 person-years, affecting an estimated 15,000 individuals in the United States and 30,000 in Europe each year (48, 49). Managing TAA presents challenges in both elective and emergency cases. Deciding on surgical treatment for elective cases is complicated and hinges on weighing the surgical risks against the potential hazard of aortic rupture. Despite advancements in medical and surgical treatments, the outcomes for emergency surgical cases, particularly regarding operative mortality and morbidity, remain relatively high (50, 51). The prevalence and incidence of TAA, the lack of effective pharmacological interventions, surgical treatment outcomes, and long-term results, regardless of initial treatment strategies, all contribute to the burden on the healthcare system for TAA patients. Identifying the underlying molecular pathways that trigger TAA onset and progression is crucial for developing therapeutic agents that can prevent aortic dilatation and dissections. In the current study, we showed that pathological NF-kB signaling in thoracic vSMCs contributes to phenotypic switching and hyperproliferation, leading to vascular remodeling in TAA. Furthermore, we discovered that many components associated with TAA progression can be reduced by inhibiting mitochondrial fission.

Mitochondrial dynamics have increasingly been linked to SMC dysfunction. Studies have shown the role of mitochondrial dynamics in vSMC phenotypic switching and proliferation (52). The protective effects of inhibiting mitochondrial fission were further supported by the suppression of PDGF-induced mitochondrial fission and vSMC migration through the expression of the dominant-negative mutant DLP1-K38A or DLP1 silencing (53). Similarly, treatment of vSMCs with PDGF showed excessive mitochondrial fragmentation, increased cell proliferation, and a transition to a synthetic phenotype, which was suppressed by mdivi-1 treatment (18). Deng et al. showed that Ang II induces hypertension and that mdivi-1 can attenuate this effect by mediating the phenotypic switching of vascular smooth muscle cells (vSMCs). Mitochondrial division inhibitor 1 (mdivi-1), which is a derivative of quinazolinone, has been reported as a selective inhibitor of mitochondrial fission (23). The mechanism of action of mdivi-1 is reported to involve the inhibition of Drp1 GTPase activity; however, the mechanistic insights into mdivi-1 are still under investigation, as studies have indicated a GTPase-independent mechanism of Drp1 inhibition (54, 55). The therapeutic effects of mdivi-1 have been noted in sepsis (25), acute myocardial infarction (24), and neurodegenerative disorders (26). The inhibition of hypoxia-triggered mitochondrial fission in vascular smooth muscle cells (vSMCs) by mdivi-1 has been reported in an ischemic/hypoxia injury rat model. This occurs through the attenuation of Drp1 activation at Ser616, with negligible effects on its GTPase activity or protein expression (21). Current literature indicates a strong association between mitochondrial dynamics and vascular dysfunction in cardiovascular diseases based on several in vitro and in vivo studies.

To clarify the protective mechanism of mitochondrial fission inhibition in TAA, we activated the renin-angiotensin system by administering Ang II to hyperlipidemic mice as a TAA preclinical model. As described previously (32), chronic administration of Ang II in ApoEKO mice led to the development of severe TAA. The subcutaneous Ang II infusion in ApoEKO mice remains one of the key research models for studying TAA pathogenesis. It displays clinical characteristics such as vascular remodeling, perivascular fibrosis, inflammatory cell infiltration, luminal expansion, and aortic dissection (5, 7, 56). The aortic tissue exhibited increased mitochondrial structural disintegration as observed through transmission electron microscopy. Immunofluorescence staining confirmed excessive mitochondrial fission linked to Ang II treatment. Treatment with a mitochondrial fission inhibitor restored the mitochondrial structure in the aortic tissue. The data confirmed the association between excessive mitochondrial fission and the onset of TAA development.

The onset and progression of TAA are characterized by pathological vascular remodeling, which involves ECM degradation and excessive collagen deposition (32, 36). The chronic administration of Ang II results in thoracic aortic dilation, accompanied by medial elastolysis and perivascular collagen deposition. Mdivi-1 has been shown to reduce collagen secretion and, consequently, fibrosis by inhibiting Drp1-dependent fission in cardiovascular pathologies (57, 58). Consistent with the literature, mdivi-1 treatment restored aortic structure and function, reducing medial disintegration and collagen deposition. vSMCs represent a primary cell type in the medial layer of the aorta and play a critical role in aortic pathologies (40). Under physiological conditions, vSMCs display a contractile phenotype and uphold vascular tone in the vessel wall. However, under pathological conditions, vSMCs may dedifferentiate into a synthetic phenotype, releasing pro-inflammatory cytokines, matrix metalloproteinases (MMPs), and collagen. (40, 59). The phenomenon of phenotypic switching often serves as a precursor to vascular pathologies. Numerous studies have linked the phenotypic switching of SMCs to a synthetic phenotype associated with vascular remodeling and aneurysm formation. (60). Consistent with our previous study (32), our results demonstrated that Ang II-induced vSMC polarization to synthetic phenotype exacerbates vascular remodeling associated with TAA development. The inhibition of mitochondrial fragmentation resulted in the restoration of vSMC contractile phenotype and attenuation of TAA. The findings were consistent with the published study, which elucidated that mdivi-1 inhibited mitochondrial fragmentation and vSMC phenotypic switching via AMPK-α downregulation in Ang II-induced hypertension (14). Increased vSMC proliferation has been linked to phenotypic switching and has been implicated in the onset and progression of TAA.

The dedifferentiation of vSMCs is characterized by increased proliferation and migration (61–63). Reduced Ki67 fluorescence in mdivi-1-treated vSMCs suggested a decrease in proliferation and a reduction in the pathological transformation associated with TAA. Mitochondria are central to cellular energy metabolism. The bioenergetics of vSMCs have been linked to mitochondrial structure and function. The phenotypic switching of vSMCs and mitochondrial fission have been associated with impaired oxidative phosphorylation (64, 65). We observed a trend of restored ATP production and increased maximal respiration, along with reduced proton leak, in vSMCs treated with mdivi-1, suggesting enhanced metabolic activity and function.

The study findings indicated a complex interplay among mitochondrial dynamics, vSMCs phenotypic switching, and hyperproliferation, contributing to the onset of TAA. Activation of the NF-kB pathway in vSMCs mediates phenotypic switching and proliferation following vascular injury (44). Drp1-dependent mitochondrial fission has been shown to enhance cell proliferation in hepatocellular carcinoma through the interaction of p53 and NF-kB pathways (66). Furthermore, mitochondrial fission was found to be a key modulator of sustained activation of NF-kB signaling in endothelial cells (67, 68). The literature suggests an interplay between mitochondrial fission and NF-kB signaling that can mediate vSMC dysfunction, leading to TAA. We demonstrated nuclear translocation of the NF-kB subunit p65 in Ang II-treated vSMCs, facilitating the transcriptional upregulation of the cell cycle kinases cyclin E1 and cyclin D1. The mdivi-1-mediated fission inhibition attenuated p65 translocation, with the downregulation of cyclins and the upregulation of the cyclin-dependent kinase inhibitor 1/p21 in vSMCs. Thus, mdivi-1 reinforced the contractile vSMC phenotype by regulating the cell cycle via NF-kB signaling.

A key limitation of this study is the use of only male mice in the preclinical thoracic aortic aneurysm (TAA) model. While TAA is more common in males, sex-specific differences in vascular biology and hormonal regulation may significantly impact disease progression and therapeutic response (69, 70). Notably, estrogen has been shown to have protective effects on vascular integrity, which could alter the impact of Ang II and mdivi-1 in female mice (69). Future studies that include both sexes are essential for providing a more comprehensive understanding of TAA pathophysiology and potential sex-specific treatment strategies. Furthermore, this study does not consider the systemic effects of Ang II and/or mdivi-1 administration in the Ang II-induced TAA model. Ang II infusion is known to cause systemic hypertension and inflammation, which may independently contribute to aneurysm formation and progression (71, 72). Furthermore, mdivi-1, a mitochondrial division inhibitor, may exert effects beyond the aorta, potentially influencing other vascular compartments or metabolic pathways (73, 74). Future research should aim to delineate these systemic effects to better isolate direct aortic mechanisms. To our knowledge, this is the first study investigating the inhibition of mitochondrial fission as a potential therapeutic target in TAA. Previously, Cooper et al. reported the therapeutic benefits of mdivi-1 in the Ang II-induced abdominal aortic aneurysm (AAA) murine model (75). The study demonstrated that Drp1-dependent mitochondrial fission plays a crucial role in mitochondrial dysfunction and the inflammatory activation of vSMCs, thus leading to AAA. In conclusion, pharmacological inhibition of Drp1-dependent mitochondrial fission reduces Ang II-induced TAA. Our study provides evidence for the potential role of mitochondrial fission in vSMCs proliferation, phenotypic switching, and metabolic dysfunction that contribute to TAA development.

## Data availability statement

The data supporting the findings of this study are available from the corresponding author upon reasonable request.

